# Morphological shifts consistent with the island syndrome in land-bridge island birds

**DOI:** 10.64898/2026.06.11.731573

**Authors:** Maarten J. K. Hoepel, Sebastian Steibl, Martim Melo, Amancio Motove Etingüe, Sonya M. Clegg, Steven C. Miller, Pau Enric Serra-Marin, Pablo Owono Nchama, Ursula Revelanda Asangono Edjang Maye, Sofía Hayden Bofill, Maximiliano Fero Meñe, Katherine Gonder, Luis Valente

**Affiliations:** Naturalis Biodiversity Center, Leiden, the Netherlands; School of Biological Sciences, University of Auckland, New Zealand; Fitzpatrick Institute, South Africa; Museum Porto, Portugal; Bioko Biodiversity Protection Program, Equatorial Guinea; Department of Biology, University of Oxford, United Kingdom; Camden County College, USA; IMEDEA (CSIC), Spain; INDEFOR-AG, Equatorial Guinea; National University of Equatorial Guinea, Equatorial Guinea; Texas A&M University, USA; Groningen Institute for Evolutionary Life Sciences, University of Groningen, the Netherlands

**Keywords:** Land-bridge islands, island syndrome, island rule, birds, subspecies

## Abstract

Land-bridge islands are former mainland areas isolated by post-glacial sea-level rise (<15,000 years) and the most common island type. Because of their recurrent connectivity with continents, it is unclear whether species on land-bridge islands can undergo evolutionary changes associated with the more isolated oceanic islands (‘island syndrome’). Here, we test the hypothesis that the selective environment on land-bridge islands exerts predictable and consistent evolutionary shifts in morphological traits of songbirds. We apply Bayesian hierarchical models to a morphological dataset of 6,917 individuals comprising 185 species of songbirds from four land-bridge islands (Bioko, Sri Lanka, Taiwan and Trinidad) and adjacent continents. Across all 185 species, we find that occurrence on a land-bridge island has clear directional effects on five morphological traits related to beak, wing, and tarsus, as well as a general increase in body size. At the species level, 57 out of 90 tested species exhibit significant morphological divergence between land-bridge island and mainland, yet for only 20 of these are the land-bridge island populations recognised as distinct endemic subspecies. Our results show that occurrence on land-bridge islands has a detectable effect on passerine morphology consistent with the island syndrome, and suggest these islands harbour previously unrecognized unique biodiversity.

## Introduction

Islands provide unique biogeographical and environmental conditions and are thus fascinating settings to study evolution (Warren et al., 2015). They are highly diverse in terms of their degree of isolation (Wallace, 1880), from remote isolated oceanic islands that were never connected to another landmass, to semi-isolated island-like environments adjacent to or within continents (e.g., “sky-islands”). Much of our understanding of island evolution processes, however, is derived from studies focusing on volcanic oceanic islands that are often species poor, but rich in endemic species (Valente et al., 2020; Whittaker et al., 2008). Oceanic island systems are devoid of species at the time of their formation, thus facilitating controlling for confounding variables when studying evolutionary processes (Barajas Barbosa et al., 2023; Whittaker et al., 2008). In contrast to oceanic islands, land-bridge islands (also known as continental shelf islands) are separated from the continent by shallow seas and were connected to the mainland in periods of low sea level (Ali, 2018). Although land-bridge islands represent over 70% of the number of islands larger than 1 km^2^ globally (Weigelt et al., 2013), they are underrepresented in evolutionary studies. This is largely due to their temporally recent isolation from, and recurrent connectivity and close proximity to the continent, resulting in biotas that resemble the surrounding continental regions, overwhelmingly composed of non-endemics and hosting higher numbers of species than oceanic islands, as well as often harbouring certain guilds (e.g., large mammals) that tend to be absent on oceanic island systems (Ali, 2018).

Despite their similarities to the continents with respect to species composition, land-bridge islands nevertheless hold fascinating species assemblages, consisting of a reduced subset of the continental species pool, and including an interesting mix of vicariant species that were present on the island before it became separated, and species that have colonized more recently via overwater dispersal (Diamond, 1972). Regardless of whether they have a vicariant or dispersal origin, the majority of species on land-bridge islands are presumed to be non-endemic – that is, they have extant conspecific populations on the continent (Hammoud et al., 2021). This is because land-bridge islands are very young from an evolutionary perspective (<15,000 years), meaning that there often has not been sufficient time for evolutionary differences to accumulate, and because potential gene flow from the nearby mainland may prevent divergence (Garg et al., 2022). However, because they are effectively in allopatry with respect to the mainland, populations of non-endemic species confined to land-bridge islands may be at the early evolutionary stages of genetic and/or morphological differentiation (Papadopoulou & Knowles, 2015). This could result in an underestimation of their biodiversity, as some land-bridge island populations may constitute independently evolving units that remain undetected, or even qualify as new subspecies that are endemic to the island. Land-bridge islands can therefore not only offer fundamental insights into the processes of early isolation among species (Itescu et al., 2020), but may also potentially harbour an underappreciated level of hidden biodiversity.

In oceanic islands, consistent evolutionary shifts in island endemics across taxa have been well-documented. These are collectively referred to as the “island syndrome”, a suite of ecological, physiological, behavioural, and morphological traits that tend to evolve in species living on islands as compared to their mainland relatives (Adler & Levins, 1994; Baeckens & Van Damme, 2020; Benítez-López et al., 2021). This macroevolutionary pattern is characterized by changes in morphology (e.g., insular dwarfism, gigantism, loss of dispersal ability), behaviour (e.g., island tameness) and life history traits (e.g., reduced litter/clutch size) (Covas, 2012; Jezierski et al., 2024; Whittaker et al., 2023). Such evolutionary changes are typically explained by release from ecological pressures due to the lack of or reduced number of species of predators and competitors on islands (Lomolino, 2016; Runemark et al., 2014; Whittaker et al., 2023). Island size and isolation, however, may impact the relative strength and effect of the abovementioned factors, generally leading to more pronounced or extreme shifts on smaller and more isolated islands compared to islands that are more similar to the continent or closer to it (Benítez-López et al., 2021). However, it remains largely untested whether land-bridge islands offer similar selective pressures conducive to consistent evolutionary changes, or whether they are simply too similar to the continent or too young for any evolutionary divergence consistent with the island syndrome to be observed.

Passerine birds provide an ideal system for addressing this topic. They are a taxonomically and ecologically diverse group that is widespread across islands around the world (Matthews et al., 2022). Ample data on traits, distribution and phylogeny of passerines is available, enabling comparative morphological analyses (Jetz et al., 2012; Jezierski et al., 2024; Pigot et al., 2020). Evolutionary changes consistent with the island syndrome are well documented in endemic passerines from oceanic islands, including an increased tendency towards reduction of flight abilities (Sayol et al., 2020; Wright et al., 2016), and for more rounded wings (Leisler & Winkler, 2015). Regarding body size, passerine species are generally small, and therefore are candidates for evolving gigantism on islands, according to the “island rule”, which states that small body-sized species become larger on islands (Andrade et al., 2015; Benítez-López et al., 2021; Clegg & Owens, 2002). Indeed, several studies have found evidence for body size increases in insular passerines, as well as for bills and legs being generally longer on islands (Bell, 2018; Clegg & Owens, 2002; Greenberg & Danner, 2013; Jezierski et al., 2024; Wright & Steadman, 2012). Whether such patterns are also observed in land-bridge island birds remains unresolved, but addressing this would constitute a powerful test of whether land-bridge islands exert similar evolutionary forces on passerines as oceanic islands. Despite their obvious dispersal prowess, studies indicate that non-migratory birds can be relatively hesitant to cross even small barriers, suggesting that the open seawater between the mainland and a land-bridge island may constitute a sufficiently strong barrier to allow island populations to become isolated (Garg et al., 2022; Harris & Reed, 2003; Johnson et al., 2017). Furthermore, dispersal is not a static trait, with studies showing evolution of reduced dispersal capacity on islands (Diamond et al., 1976; Estandía et al., 2023). Consistent with the idea that the sea between the land-bridge and the mainland poses a dispersal barrier, studies of land-bridge island bird populations have shown indications of divergence in both morphology and genetics (Burridge et al., 2013; Chen et al., 2015; Dong et al., 2020; Dudaniec et al., 2011; Marcaigh et al., 2021; Wright & Steadman, 2012). However, previous studies have centred on individual species and islands, and therefore the extent to which land-bridge island populations generally conform to the island syndrome remains unknown. Here we focus on the morphological aspects of the island syndrome, particularly on body size shifts related to the island rule, but we stress that the suite of traits included in the island syndrome is much broader (e.g., including life-history, physiological and behavioural traits).

Beyond potential predictable morphological shifts consistent with the island syndrome, neutral or random processes occurring during and after the establishment of isolated populations (e.g., founder effects, genetic drift) may drive individual, species-specific changes in morphology (Anderson & Weir, 2022; Nosil & Flaxman, 2011; Spurgin et al., 2014). Therefore, within land-bridge islands, we may expect to find passerine lineages in the early stages of divergence, some of them possibly warranting the treatment as unique endemic subspecies. Nevertheless, because land-bridge island populations of passerine birds are generally perceived to be part of the mainland meta-population, they are often not considered separately in macroecological or macroevolutionary analyses and can be overlooked in conservation studies.

Here, we address these research gaps by conducting a comparative morphological analysis of multiple passerine species across four different land-bridge islands. We integrate morphological data from field and museum specimens on ten morphological traits across 185 land-bridge island passerine species, comparing island and mainland populations. Our aims are twofold: 1) to identify whether there are consistent shifts in morphological traits across multiple species of passerines on land-bridge islands, and whether these shifts are consistent with expectations from the island syndrome regarding morphology; 2) to identify specific species that have undergone significant morphological divergence on land-bridge islands and may warrant treatment as subspecies. Regarding our first aim, we hypothesize that passerines on land-bridge islands display morphological trait shifts in line with island syndrome predictions, including an increased body size, beak, and tarsus length (as predicted by the island rule for species that are small on the mainland), and shortened wings and hand-wing index (linked to dispersal ability). Regarding our second aim, we expect that predictable selective pressures in combination with random genetic drift will lead to many land-bridge island populations being morphologically differentiated, which might warrant a rethinking of their taxonomic (and potentially conservation) status.

### Methodology

#### Study system and taxon selection

We focus on four geographical systems consisting of a land-bridge island and the adjacent mainland: Bioko, Trinidad, and Sri Lanka in the tropics, and Taiwan in the subtropics (Figure 1). We selected these regions based on their geographical spread, variation in species composition and data availability. We refer to these regions by the name of the land-bridge islands.

**Figure 1.**
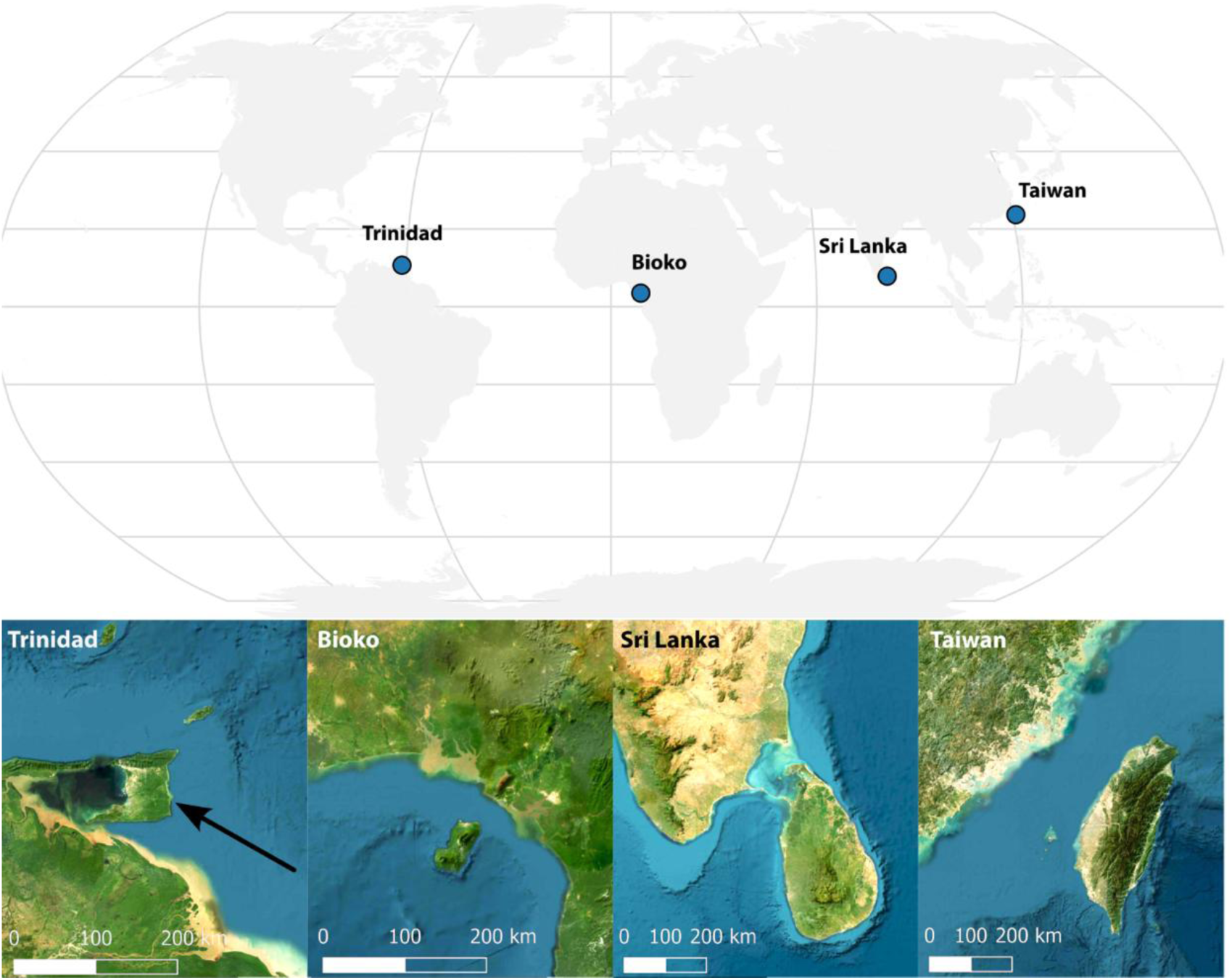
The four land-bridge island systems that are the focus of this study: Trinidad (South America), 4,769 km^2^; Bioko (West Africa), 2,017 km^2^; Sri Lanka (South Asia), 65,208 km^2^; Taiwan (East Asia), 34,507 km^2^. Insets show satellite imagery (Esri World Imagery) illustrating each island’s proximity to the mainland and position on the continental shelf.

From each of these regions we sampled native resident passerine bird species. We excluded species that a) are island endemics and therefore lack mainland populations, b) are migratory, nomadic, or vagrant; or c) have an unresolved taxonomy, e.g., making it unclear how to attribute specimens to specific species. Our final species list depended on the availability of data, particularly whether multiple specimens from island and mainland were available. These assessments were based on the species entries in Birds of the World (Billerman et al., 2020) and the species-wide data availability from AVONET (Tobias et al., 2022).

#### Trait data

We compiled a dataset of 10 standardized morphological trait measurements derived from the AVONET database (Tobias et al., 2022), which provide extensive information on a bird’s ecomorphology: (1) beak length culmen (measured from tip to skull along the culmen), (2) beak length nares (measured from tip to the anterior edge of the nares), (3) beak depth, (4) beak width, (5) tarsus length, (6) primary wing length (measured from the carpal joint to the wingtip on the unflattened wing), (7) secondary wing length (measured from the carpal joint to the tip of the outermost secondary feather) (8) Kipp’s distance, (9) tail length and (10) hand-wing index. Eight of these are direct measurements. Kipp’s distance is calculated as the difference between the primary wing length and the secondary wing length. The hand-wing index is the fraction of the Kipp’s distance to the primary wing length (Claramunt, 2021).

We obtained trait measurements for these 10 traits based on three sources: 1) the open-access dataset AVONET, which contains data from specimens measured in the field (which will be referred to as “Field AVONET”) and measured from museum collections (“Museum AVONET”) (Tobias et al., 2022). 2) new field data collected for this study on the land-bridge island of Bioko (“Field Bioko”) and 3) new data from museum specimens from the xx museum (removed to allow double-blind review) measured for this study (“Museum xx”).

Morphological field measurements in Bioko were performed by xx. in field expeditions in 2020 and 2022. The new museum measurements were performed by xx in the period from November 2024 to February 2025, following the protocol described in the supplementary material of Tobias et al., (2022), which was based on previous studies (Borras et al., 2000; Clark, 1995). We tested consistency of measurements (individual error) by remeasuring the direct measurements for 23 specimens, in which 95% of the repeated measurements fell between a 6% decrease and 8% increase compared to the original values (Table S2). We deemed this sufficiently consistent to incorporate the data in the large dataset.

We filtered the dataset to remove all juvenile specimens, all specimens with an unknown country of origin, and all oceanic island specimens, yielding a dataset of adult individuals from both mainland and land-bridge islands (Table S1).

#### Principal component analysis of all species

We used R version 4.5.1 and RStudio (version 2025.05.1 Build 513) for our statistical analyses. We used the R packages *brms* (Bürkner, 2017) and *loo* (Vehtari et al., 2024) for the Bayesian analysis. We used *ggplot2* (Wickham, 2016), *cowplot* (Wilke, 2015), *ggdist* (Kay, 2023) and *tidybayes* (Kay, 2020) for data visualisation. To create the map in figure 1 we used *sf* (Pebesma & Bivand, 2023) and *rnaturalearth* (Massicotte & South, 2017) and *ggspatial* (Dunnington, 2017). All the code is stored in the folder “Supplementary file (code).”

To identify the main axes of morphological variation present in our dataset we performed a principal component analysis (PCA), selecting the five traits with the highest trait coverage (Table S3). These were: beak length, beak depth, tarsus length, primary wing length, and tail length. All traits were log-transformed before PCA. In total, data was available for all these five traits for 3,301 specimens, and these were included in the PCA. We selected the first two principal components for our further analyses, which together explained 78.5% of the variance.

#### Passerine-wide model

To investigate whether and how morphological traits shift on land-bridge islands (aim 1), we used Bayesian hierarchical models (Bürkner, 2017). The goal of these analyses was to test whether presence on a land-bridge island has an effect on morphology across the entire passerine dataset including multiple species and land-bridge islands. We chose Bayesian hierarchical models as they are well-suited to handle the multiple sources of variation in our dataset, as well as imbalances in sample sizes, including low sample sizes for some species.

We fitted models separately to seven of the ten morphological traits and to the first two principal components. We excluded three traits: beak length nares, secondary wing length and Kipp’s distance. Beak length nares was excluded due to high collinearity with beak length culmen (Spearman’s ρ = 0.95). Secondary wing length and Kipp’s distance were excluded because of their high collinearity with primary wing length (Spearman’s ρ = 0.95) and the hand-wing index (Spearman’s ρ = 0.85), respectively. This collinearity arises from both these measurements being used to calculate the hand-wing index.

For each trait, we selected the preferred Bayesian model by comparing a set of candidate models varying in group-level effects. We compared the candidate models using both leave-one-out cross-validation and the widely applicable information criterion (WAIC) with R package *loo* (Vehtari et al., 2024). We considered models to have the same expected predictive accuracy if the difference between the expected log pointwise predictive density was small compared to its standard error (Vehtari et al., 2017). Under this approach, we favoured the model that had the best balance between explanatory power and simplicity. In total, we fitted nine models for each of the following traits: beak length culmen (which will be referred to as beak length), beak depth, beak width, tarsus length, primary wing length, hand-wing index, tail length, principal component 1 (PC1) and principal component 2 (PC2). All candidate models for each trait are described in Table S4.

The best-balanced model was found to be the same across all traits. That model is described here, where *Y*_*ij*_ is the observed trait value of species *i* and data type *j*, which is assumed to follow a normal distribution 𝒩 with expected mean trait *Y^*_*ij*_ and standard deviation

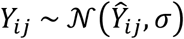

We modelled the expected mean trait *Y^*_*ij*_ as:

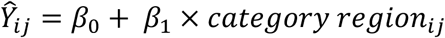

Here, β_0_ is the global intercept, β_1_ is the main effect of *category region* (either continent or land-bridge island), *i* is a group-level effect for *species* capturing phylogenetic variation, and *j* is a group-level effect for *data type* to account for methodological variation among field and museum sources. The latter also corrects for the observation that museum specimens might have lower beak and tarsus measurements due to post-mortem shrinkage and therefore a bias might arise when comparing museum data with live specimens (Wilson & McCracken, 2008). A more detailed description of each element in the Bayesian models is shown in Table S5.

For each morphological trait, we fitted the same model with four independent MCMC chains. Morphological traits were log-transformed. We also ran analyses with the two main principal components as explanatory variables. These were not log-transformed, as they already showed a gaussian distribution. For each trait model, we selected weakly informative priors and different model settings to introduce moderate model constraints to stabilise MCMC sampling (Table S6-S7). A prior sensitivity analysis was used to evaluate how strongly the model estimates were based on the choice of priors, ensuring results were driven by data rather than prior assumptions (Depaoli & Van De Schoot, 2017). We assessed the MCMC sampling and model convergence using visual inspections (trace plots), Gelman and Geweke diagnostics, R-hat statistics (all Ȓ < 1.01) and effective sample sizes (ESS > 1000). We report the model estimates by using 95% posterior credible intervals on a log-scale.

We evaluated the hypothesis that presence on a land-bridge island has an effect on a given trait or combination of traits (PC1 and PC2) by computing the posterior probability that the effect of land-bridge island on that variable was higher or lower than zero. To ease interpretation of the posterior means in Figure 2, we back-transformed posterior estimates to the original scale.

**Figure 2.**
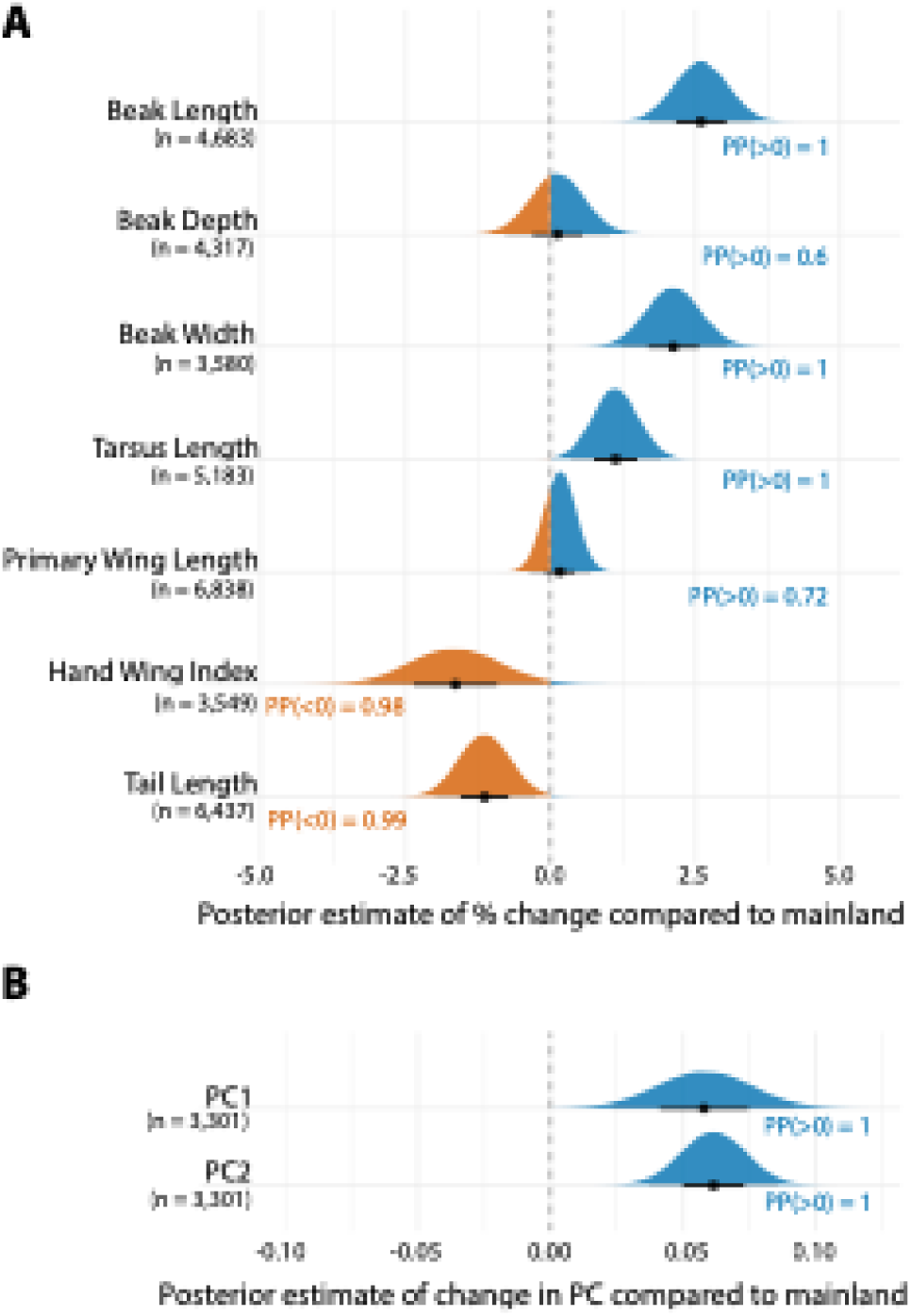
Models of morphological change trends in passerines on land-bridge islands (passerine-wide models) (A) Posterior estimates of percental change in seven morphological traits compared to the continent. Blue and orange indicate respectively positive and negative effects with the posterior probability of divergence shown next to the plot. (B) Posterior estimates of divergence for the first two principal components including five morphological traits. PC1 mostly represents body size, and PC2 beak shape. A total of 185 species are included in these analyses. Dashed line shows no difference with the continent and horizontal bars indicate 66% and 95% quantile credible intervals. PP <0 or PP >0 indicates the posterior probability that the estimated effect differs from zero, and thus differs from the mainland value.

Our final preferred Bayesian hierarchical models account for species-level variation using a group-level intercept for species. However, species are not equidistant and phylogenetic shared common ancestry can confound the results of morphological comparative analyses across species – for instance, if taxa from the same genus or family show strong effects of presence on land-bridge island because of a given characteristic that is unique to that genus or family. While we do not model phylogenetic covariance explicitly, we investigated for a possible effect of phylogenetic relationships by fitting a set of candidate models adding genus or family as nested group-level effects (Table S4). Those models did not improve explanatory power and were therefore not favoured, suggesting that higher level phylogenetic structure does not meaningfully impact our results. Thus, we are confident that our model sufficiently captures the phylogenetic variation relevant to our system.

#### Species-specific models

The passerine-wide analyses described above allow us to investigate general trends across multiple passerine species for each trait at a time, but do not inform us on the way each species may be varying between island and mainland when multiple traits are considered. To assess for each species whether land-bridge island populations are morphologically differentiated from mainland populations (aim 2), we performed a species-specific PCA and a subsequent Bayesian multivariate model. We selected the same five traits that we used in our earlier PCA on the whole dataset: beak length, beak depth, tarsus length, primary wing length and tail length. In the passerine-wide model the group-level effect “data type” sufficiently captured differences between datasets. In contrast, for some per-species comparisons, preliminary analysis showed that field data from the source “Field Bioko” was incompatible with data from other studies. This led us to remove the “Field Bioko” specimens (n = 437) from species-specific analyses. To ensure sufficient specimens were included per species, we removed all species with fewer than four specimens each from both the mainland and the land-bridge island. This resulted in 90 species for the species-specific models, including 4 species from Bioko, 11 from Taiwan, 28 from Sri Lanka and 47 from Trinidad.

We performed a PCA on each of these 90 species and kept the first two components, which together captured the majority of trait variation. These two components were then analysed in a Bayesian framework. Here, let *PC*1_*j*_ and *PC*2_*j*_ indicate the scores of individual *j* from any of the species with 𝒩 as normal distribution and σ represents standard deviation. These were jointly analysed in the multivariate model:

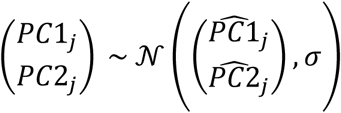

For each component, the expected value was modelled as a function of category region (either mainland or land-bridge island):

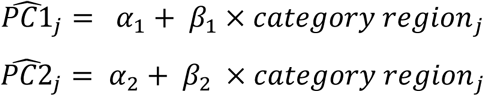

Here, α_1_ and α_2_ are the global intercepts for PC1 and PC2 respectively, β_1_ and β_2_ are the main effects of *category region* (either mainland or land-bridge island) on each of the PCs. We used this to obtain the posterior distribution of the effect of presence on a land-bridge island on both PC1 and PC2. For each principal component, we tested whether the effect was greater or less than zero, allowing us to identify species for which there is a high probability of divergence between island mainland for the individual principal components. Alongside the model estimates, we also stored the PCA loadings for each species to identify which particular traits might drive morphological divergence for that particular species.

We categorized each of the 90 species based on whether they differ between island and mainland across the five traits. We classified them into three different categories according to the level of differentiation: high posterior probability of divergence (PP(β < | > 0) > 0.95) for both PC1 and PC2 (category 1), only for PC1 or for PC2 (category 2), and for neither of the PCs (category 3, Figure 4).

Finally, for all 90 species, we checked whether the land-bridge island population is considered an endemic subspecies to that island according to Birds of the World (i.e., which taxa are considered to have endemic subspecies in Bioko, Sri Lanka, Taiwan or Trinidad) (Billerman et al., 2020). We then counted how many species in each PC differentiation category have recognised endemic subspecies on their respective land-bridge island. We would expect that species in differentiation category 1 and 2 would be more likely to be considered endemic land-bridge island subspecies.

## Results

### Dataset characteristics

Our dataset (Table S1) contains measurements for 10 morphological traits for 6,917 specimens, belonging to 185 passerine species with a median of 19 specimens per species, and a mean of 37 specimens per species (min = 2, max = 553). A total of 148 genera and 49 passerine families are represented. Species were divided into four regions, based on the land-bridge island the species was found on. For Bioko, there are 2,806 specimens belonging to 57 species; for Sri Lanka, 463 specimens belonging to 29 species; for Taiwan, 260 specimens belonging to 18 species; for Trinidad, 3,388 specimens belonging to 81 species. Our data was divided into four categories, based on the sources: “Field AVONET” (n = 2,754), “Field Bioko” (n = 437), “Museum AVONET” (n = 2,506) and “Museum xx” (n = 1,220). The dataset includes 1,392 specimens from land-bridge islands and 5,525 from the mainland. Summary statistics per trait are shown in Table S3. The over 1,500 new measurements conducted for this study (“Field Bioko” and “Museum xx”) allowed us to add a total of 119 species that would otherwise not have sufficient data in AVONET to run the analyses.

### Principal component analysis of all species

We performed a PCA on all individuals for which complete measurements for five traits (beak length, beak depth, tarsus length, primary wing length and tail length) were available (Figure S1a). All 185 species were represented. The first principal component (PC1) explained 66.5% of the variance and had approximately equal loadings (in terms of weight and direction) for each trait (Table 1). We therefore treat PC1 as being a good proxy for body size.

**Table 1.**
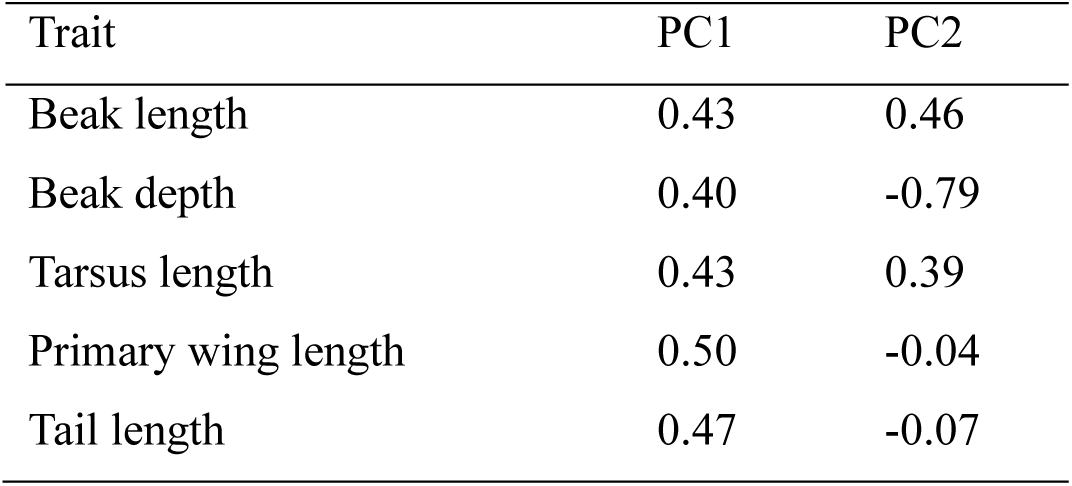
Loadings of each trait for PC1 and PC2.

The second principal component (PC2) explained 12% of the variance and was mostly driven by beak shape (positive loading for beak length and negative for beak depth) and to a lesser extent by tarsus length, and thus can be seen as being generally related to shape. To show the range of trait variation in our dataset, we highlight four indicative species in the PCA figure (Figure S1b).

### Passerine-wide models

We found strong support for our hypothesis that occurrence on land-bridge islands has an effect on morphological traits across species (Figure 2a). We found a positive effect of presence on a land-bridge island on beak length (posterior mean noted as β_1_ = 0.03 [95% credible interval of the posterior: 0.02 – 0.04], posterior probability PP(β_1_ > 0) = 1), beak width (β_1_ = 0.02 [0.01 – 0.03], PP(β_1_ > 0) = 1) and tarsus length (β_1_ = 0.01 [0.00 – 0.02], PP(β_1_ > 0) = 1). We found strong support for occurrence on land-bridge islands having a negative effect on hand-wing index (β_1_ = -0.02 [-0.03 – 0.00], PP(β_1_ < 0) = 0.98) and tail length (β_1_ = -0.01 [-0.02 – 0.00], PP(β_1_ < 0) = 0.99). We found no evidence that occurrence on land-bridge islands leads to a shift in beak depth (β_1_ = 0.00 [-0.01 – 0.01], PP(β_1_ > 0) = 0.6) and primary wing length (β_1_ = 0.00 [0.00 – 0.01], PP(β_1_ > 0) = 0.72). We fitted the same model to PC1 (proxy for body size) and PC2 (beak morphology). The model provided strong support for the hypothesis that occurrence on land-bridge islands leads to positive shifts for both PC1 (β_1_ = 0.06 [0.02 – 0.09], PP(β_1_ > 0) = 1) and PC2 (β_1_ = 0.06 [0.04 – 0.09], PP(β_1_ > 0) = 1) (Figure 2b).

### Species-specific models

We used Bayesian models to test the hypothesis that occurrence on a land-bridge island leads to shifts in PC1 and PC2 for 90 different passerine bird species (Figure 3). We found strong evidence of divergence for both PC1 and PC2 in five species (category 1): *Chlorophanes spiza*, *Dysithamnus mentalis*, *Myiodynastes maculatus*, *Myrmotherula axillaris* and *Tolmomyias sulphurescens* (Figure 3, 4; Figure S2; Table S8). These five species are all from Trinidad. We further identified 52 species for which there is strong evidence of divergence for either only PC1 or only PC2 (category 2, Figure 3, 4; Figure S3, S4; Table S8). For 33 species, there was no evidence that occurrence on a land-bridge island has a directional effect on the trait space (category 3, Figure 3, 4; Figure S5; Table S8). The percentage of variance in each species explained by PC1 ranged from 33.9% to 67.5%, and for PC2 from 16.1% to 35.1%.

**Figure 3.**
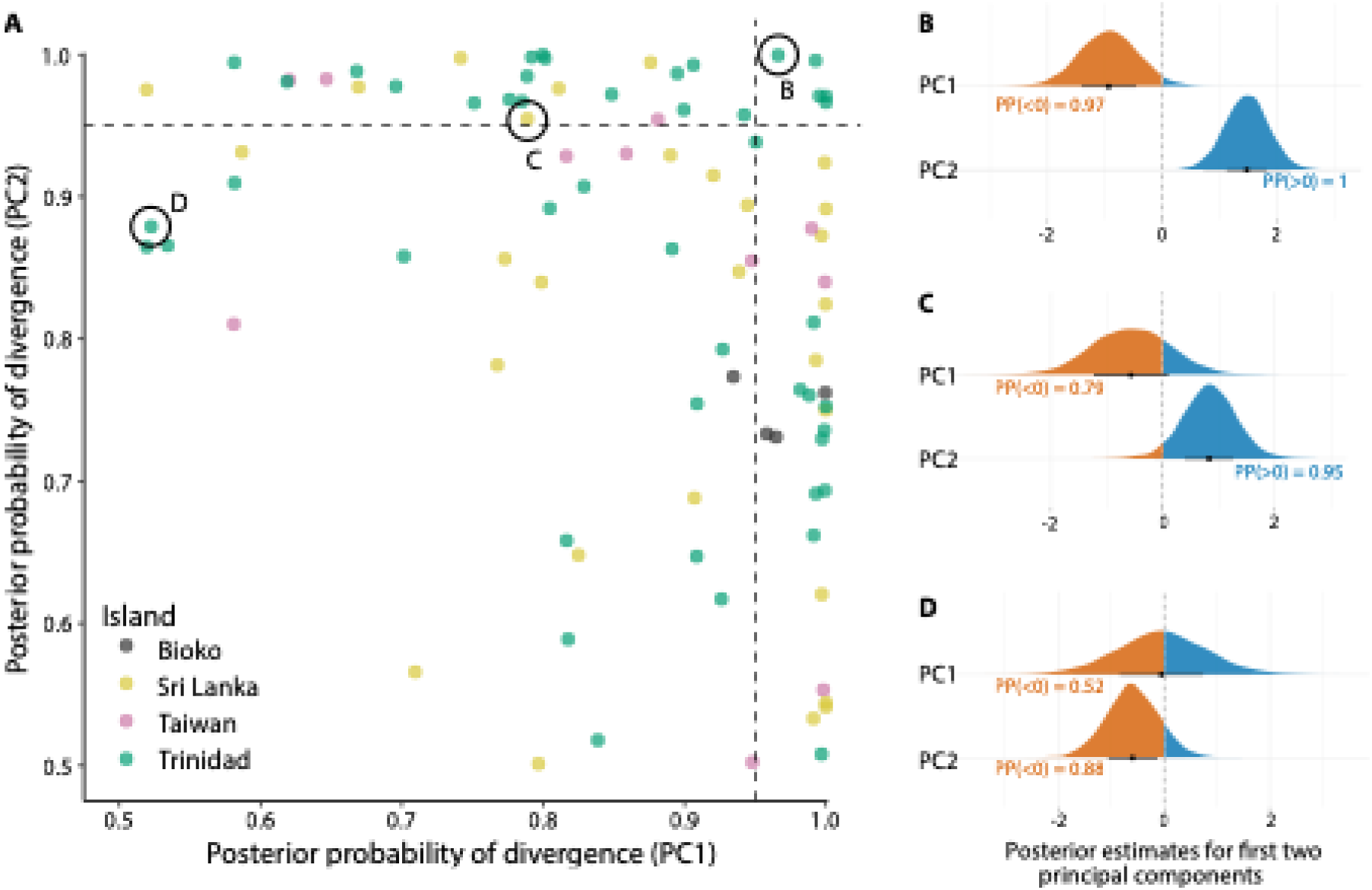
Summary of the results of the models of morphological divergence of passerine birds on land-bridge islands for 90 species (species-specific models) A) Scatterplot indicating the mean posterior probabilities of morphological divergence between land-bridge island and mainland populations across species. Each point represents a species and shows the posterior probability that the effect of category region differs from zero for the first (PC1) and second (PC2) principal components. Colours indicate the land-bridge island (Bioko, Sri Lanka, Taiwan, Trinidad). Dashed lines mark the 0.95 threshold used to classify a component as showing a high probability of divergence. Panels on the right (B-D) show posterior estimations of the effect for PC1 and PC2 (orange is negative and blue is positive) for three example species representing each category of divergence: B) *Chlorophanes spiza* (category 1, divergence in both PCs), C) *Chloropsis jerdoni* (category 2, divergence in one of the PCs) and D) *Tyrannus melancholicus* (category 3, no divergence in either PC). The three species are highlighted in A with a bubble. Detailed results for each species are given in Table S8.

**Figure 4.**
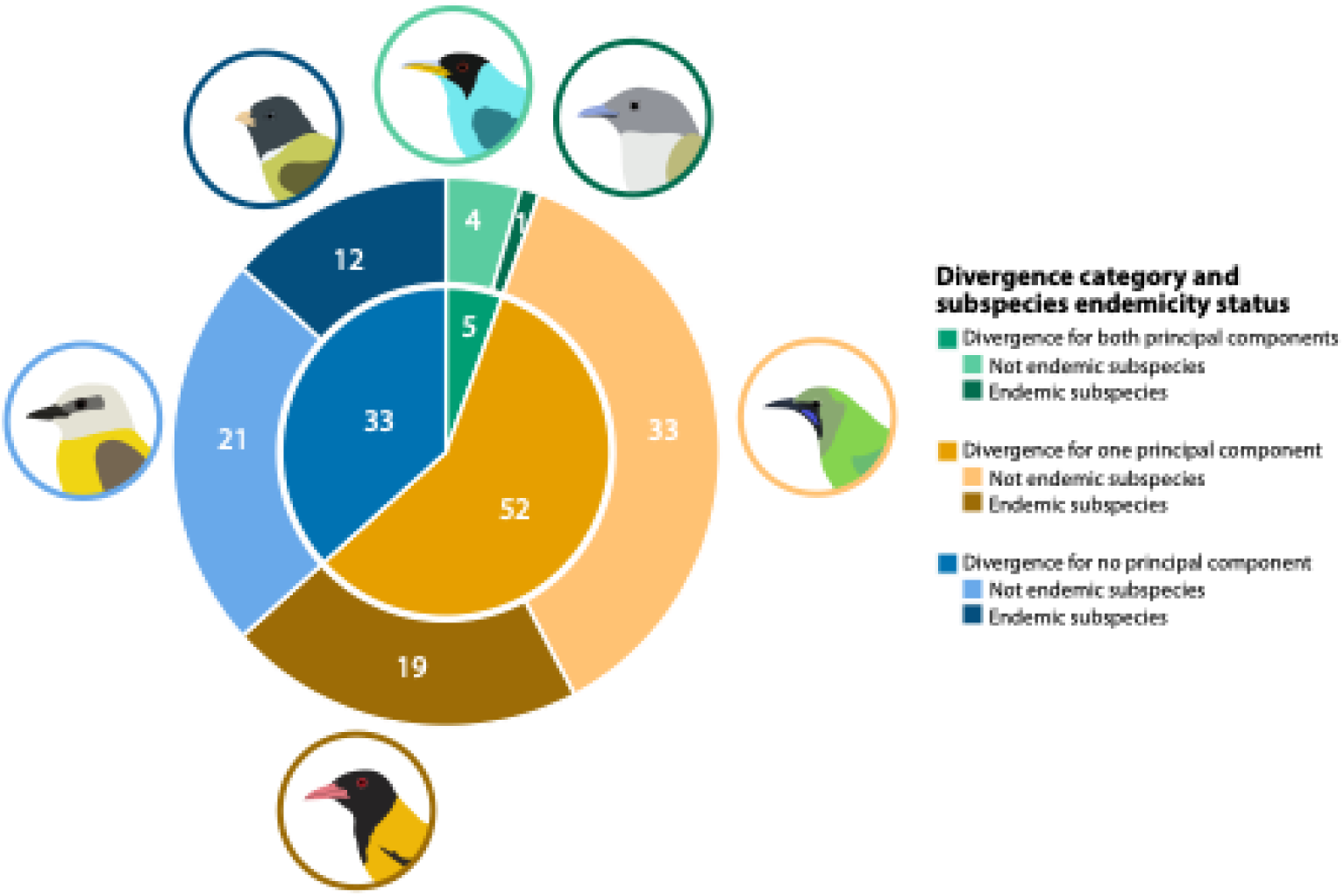
Classification of the 90 taxa included in the species-specific models according to divergence category and subspecies endemicity status. The inner circle of the donut chart shows the number of species in each of the three categories of divergence based on the results of the Bayesian models: divergence for both principal components (category 1, green), for one principal component (category 2, orange) and for no principal component (category 3, blue). The outer ring separates those species into whether they have land-bridge island populations currently considered as endemic subspecies to the land-bridge island (according to Birds of the World). Darker shade – land-bridge island population considered endemic subspecies; lighter shade – land-bridge island population not considered endemic subspecies. Species illustrated are, clockwise starting from light-green at the top: *Chlorophanes spiza, Tolmomyias sulphurescens, Chloropsis jerdoni, Oriolus xanthornus, Tyrannus melancholicus and Spizixos semitorques*.

Of the five species that had high posterior probability of divergence for both PC1 and PC2 (category 1), only one, the *Tolmomyias sulphurescens*, is currently classified as a land-bridge island endemic subspecies according to Birds of the World (Billerman et al., 2020). For the 52 species with high posterior probability of divergence for either one of the principal components (category 2), 19 are currently defined as a land-bridge island endemic subspecies (Figure 4, Table S8). For the 33 species with inconclusive evidence of divergence (category 3), 12 are classified as a land-bridge island endemics subspecies.

## Discussion

Our analyses of morphological traits of passerine birds revealed surprisingly clear trends of morphological evolution in land-bridge island populations compared to the mainland. In general, the trends we detected are compatible with the early evolution of the island syndrome and consistent with the island rule. In addition, we found that for many species (63%) there is some degree of morphological differentiation of land-bridge island populations compared to the mainland. Below, we interpret these results considering the dynamic biogeography of land-bridge islands.

### Trait shifts: body size, beak, tarsus, tail

We found small but consistent changes in several morphological traits for passerines on land-bridge island compared to the continent, suggesting predictable evolutionary shifts in songbird morphology under the selective pressures found on land-bridge islands. Across the 185 species we found an increase in PC1, which is proxy of a bird’s overall body size (Table 1 and Bell, 2018), on land-bridge islands. Such an increase in overall body size fits the island rule, which proposes that species converge on a body size optimum owing to multiple ecological drivers (e.g. resource limitation, limited predation). This means that smaller species, which passerines generally are, will usually increase in size on islands (Benítez-López et al., 2021; Clegg et al., 2002; Jezierski et al., 2024). In the traits we measured, there were small but significant increases in overall size for the beak and the tarsus.

For beak-related measurements, we found increases in beak length (∼3% change) and beak width (∼2% change), but no estimated shift in beak depth. Combined with the observed effect of land-bridge island presence on PC2, which is strongly linked to bill morphology, this suggests that bird individuals on land-bridge islands tend to have longer, slender beaks compared to mainland individuals, but not necessarily deeper beaks. This fits the long-standing observation that island passerines tend to have larger beaks than their mainland relatives (Grant, 1965; Jezierski et al., 2024). Beaks are linked to a wide range of functions, such as thermoregulation, nest construction, and of course dietary functions (Navalón et al., 2019). An increase in beak length can be driven by niche broadening, allowing a bird to forage for a broader selection of food, usually because larger food items can now be consumed (Grant, 1965; Wright & Steadman, 2012). This generalist behaviour has previously been described for oceanic islands (Clegg & Owens, 2002), and also documented before for land-bridge island passerines (Keeler-Wolf, 1986). However, individual-level studies of island birds have found cases where generalist populations consist of different types of individual specialists, suggesting that morphological shifts may not necessarily be explained by niche broadening and generalist foraging (Scott et al., 2003).

We found a positive, yet small change in tarsus length (∼ 1 %) on land-bridge islands. An increase in tarsus length might be explained by two different processes. Larger tarsi are associated with using a broader range of perch types (Carrascal et al., 1994), as well as ground foraging and the adoption of an increased terrestrial lifestyle (Ponti et al., 2025). This may further contribute to the previously mentioned niche expansion (Grant, 1965; Wright et al., 2016). Another hypothesis is related to reduced dispersal abilities. As legs initiate the flight take-off using a leg-thrust (Earls, 2000), longer legs can compensate in flight initiation for birds with reduced dispersal abilities (Wright et al., 2016). Both explanations might be applicable to different species and disentangling these related ecological processes remains a challenge for future studies.

Finally, while general body size (PC1) increases, we observed a small decrease in tail length (∼ 1% change) for land-bridge island specimens. Tail length is often associated with flight optimization, where longer tails are aerodynamically optimal (Balmford et al., 1993). Additionally, tail elongation is generally attributed to sexual selection (Fitzpatrick, 1999). In this context, a reduction in tail length could be expected given the previously discussed potential decrease in dispersal ability, as well as the general pattern of reduced ornamental traits (e.g. ‘duller’ plumage) for island birds (Doutrelant et al., 2016; Jezierski et al., 2024). However, both the aerodynamic and the ornamental values depend on more than just length, as they are also dependant on shape (e.g., presence of a fork) and width (Dawideit et al., 2009). These measurements are currently unavailable for the assessed species and any interpretation of the observed reduction in tail length should be done with caution.

### Trait shifts: wing shape and dispersibility

Wing-related traits are often linked to dispersal ability and should be especially relevant for explaining avian distributions on islands. The hand-wing index in particular is generally considered to be a good predictor of dispersal ability (Claramunt, 2021; Claramunt et al., 2022). Accordingly, we found some evidence for reduced dispersal abilities in our data as we observe a posterior estimation of a general trend towards a decrease in the hand-wing index on land-bridge islands (∼ 2% change). The reduction in flight-ability in birds on islands is a widely observed phenomenon (Sayol et al., 2020). Due to a reduction in predation risks, flying becomes less necessary, and as it is energetically consuming, the flight apparatus eventually becomes redundant (Jezierski et al., 2024). A reduction in hand-wing index should therefore be an indication of this process among island-dwelling passerines. However, while an overall reduction in wing size is associated with poorer dispersal ability (Wright et al., 2016), we did not observe a change in primary wing length in our models, suggesting that the reduction in hand-wing index is primarily linked to a change in the shape of the wing (i.e., more rounded) rather than a reduction of size.

While the reduction in hand-wing index and potentially associated reduction in dispersal ability seems to align well with island syndrome expectations (Jezierski et al., 2024), the relationship between insularity and dispersal ability is complex, particularly for land-bridge islands. For bird species capable of readily crossing open water, the land-bridge island may form part of a meta-population. However, for less dispersive species, such as understory forest birds, the picture may be very different, and their presence on the island may derive from the periods when it was part of the continent (Diamond, 1972). To add to this complexity, dispersibility itself evolves, and species that readily traverse large distances on a mainland area might not necessarily cross water barriers, such that population differentiation builds more quickly in true island situations, even when direct distances are small (Estandía et al., 2025; Radu et al., 2024). Interestingly, studies have shown that islands are in fact positively associated with an overall higher hand-wing index, particularly more isolated oceanic islands (Bastidas-Urrutia et al., 2025; Sheard et al., 2020). A higher hand-wing index (presumably linked to higher dispersibility) may be necessary to allow species to reach oceanic islands, potentially obscuring any effect of loss of dispersibility. In contrast, and due to the former land connection, this filter probably does not exist for land-bridge islands (Bastidas-Urrutia et al., 2025) so the pressure for higher hand-wing index exerted on colonisers may never kick in for vicariant land-bridge island taxa. Alternatively, a selective pressure that may be present is one to become more sedentary, as individuals attempting to leave the island would encounter the recently-formed water barrier between the island and the continent. In addition, the selective pressure for escape flights may perhaps become reduced if land-bridge islands see a reduction in predation pressure, for instance associated with extinction of predators following biodiversity relaxation after the island becomes separated from the mainland (Diamond, 1972).

### Magnitude of shifts

The ecological and evolutionary setting of land-bridge islands lies somewhere between insularity and continentality. Throughout their history, they have been connected and disconnected multiple times, and therefore have unique characteristics of islands (e.g., isolation, small area) but also many aspects that resemble the continent (e.g., higher species diversity than oceanic islands, low endemism). When studying shifts in traits on land-bridge islands across multiple passerine species, we assumed that they have similarities with oceanic islands in terms of changes in selective pressure (e.g., ecological release, predator relaxation), but to a lesser extent. In other words, selective pressures typical of islands may be present (e.g. lower diversity of competitors and of predators) but without complete insularity being achieved (Ali, 2018). This contrasts with classic oceanic islands, which tend to have highly disharmonic faunas and floras. In that sense, the relatively small shifts in traits between 1-3% change supported by our results are to be expected. However, while this percentage difference may seem small, owing to allometry it may actually lead to very large changes in body size (Van Der Geer et al., 2018). Furthermore, given that a key premise of the evolution of island syndrome traits is the absence of or reduced number of predators and competitors (Lomolino, 2005), it could be seen as surprising that we found such consistent shifts at all, because a diverse community of species is present on all the land-bridge islands we studied, including large carnivorans and large mammalian herbivores.

Our results may underestimate the magnitude of evolutionary trends on land-bridge islands because we include species that may vary strongly in their degree of isolation – e.g., some may be truly isolated with no gene flow with the mainland, whereas others may experience ongoing or recent gene flow. Due to the broad sampling approach of this study, we were unable to include genetic data. Future studies including population genetic data could quantify the degree of genetic isolation and estimate the amount of time a given species has been present on the islands. Molecular data would also help identify whether species were of dispersal or vicariant origin on islands (Burridge et al., 2013). Sustained gene flow at sufficient rates between land-bridge island and mainland populations would likely counterbalance any of the observed shifts. Therefore, identification and removal of well-connected species from our dataset would likely result in detection of more pronounced trait shifts, especially for vicariant species where the island populations became isolated in earlier glacial cycles. While here we focused on morphological aspects of the island syndrome, in the future it would be interesting to investigate and collect data on further components of the syndrome - behavioural, physiological, life history - to determine if these components show a similar pattern of expected direction but small magnitude of shifts in land-bridge islands.

### Species-specific shifts: consequences for taxonomy and conservation

Following a period of isolation, island populations should become morphologically differentiated either due to predictable trait shifts associated with local insular selective pressures, or due to stochastic or neutral processes shaping morphology in allopatry (Anderson & Weir, 2022). Thus, regardless of the classic ecological features of insularity, isolation between any two populations should be conducive to the evolution of measurable morphological changes (Anderson & Weir, 2022). To examine whether land-bridge islands populations follow these patterns, we explored species-specific morphological divergence in more detail.

For 57 out of 90 species included in the species-specific analyses (63%), we found significant morphological divergence for one or both of the principal axes of intraspecific variation between land-bridge and continental individuals. This suggests some degree of genetic isolation between island and mainland populations for the majority of species we analysed. In contrast, for 33 species (37%), we did not find significant morphological differentiation divergence for any of the principal components. For these species, it may be that divergence is too recent (e.g., recent colonisers), that gene flow is ongoing, or that the differences in selective environment between island and mainland have not (yet) favoured morphological differentiation.

We checked subspecies designation in the 90 species we examined, and found that a total of 32 species have their land-bridge island populations currently classified as endemic subspecies according to Birds of the World (Billerman et al., 2020) (Table S8). Interestingly, there was no clear correspondence between subspecies classification and the degree of divergence in the morphological traits we considered. Out of the 57 species that show some degree of differentiation, only 20 have recognised endemic subspecies on the land-bridge island. This suggests that there may be some currently unrecognised subspecies-level diversity on land-bridge islands worldwide. In contrast, for 12 land-bridge island populations currently treated as distinct subspecies, we found no evidence of differentiation in the five traits we analysed. These are also good candidates for clarification of their taxonomic status using multiple lines of evidence.

Defining and delimiting subspecies is notoriously difficult and even subjective, but is an useful exercise towards labelling independent evolutionary units for conservation prioritization, such as in the “Evolutionary Significant Units” framework (Moritz, 1994). This study provides a strong indication that there may indeed be evolutionary unique land-bridge island populations deserving of particular conservation attention. Island species are especially vulnerable to anthropogenic activities (Fernández-Palacios et al., 2021; Matthews et al., 2024), and land-bridge islands may harbour unique island-adapted populations that merit tailored conservation action. Subspecies endemic to islands face similar threats as endemic species (Szabo et al., 2012) and, therefore, discerning the taxonomic status of island populations may have tangible conservation value (Phillimore & Owens, 2006). Further studies incorporating evidence of genetic, behavioural, or song divergence may help ascertain whether these land-bridge populations should indeed be given a subspecies status. This will allow us to identify particularly isolated land-bridge island passerine populations that may constitute cryptic biodiversity.

Finally, out of the 57 species showing some morphological differentiation, only five exhibited divergence in both principal components, all five from the island of Trinidad. The exclusion of endemic species from our dataset led to the removal of island species with a high degree of morphological divergence compared to the mainland. At the same time, this filtering step reduced the total amount of species available per land-bridge island, particularly for Taiwan and Sri Lanka, which have higher numbers of endemic bird species (32 and 35, respectively) compared to Trinidad and Bioko (1 and 2, respectively) (Lepage et al., 2014). Taiwan and Sri Lanka are both larger islands and offer diverse habitats, allowing for a higher chance of speciation (Kisel & Barraclough, 2010), resulting in the higher numbers of endemics (Whittaker et al., 2023). This difference in number of endemic species per island resulted in an imbalance in the origin composition of our dataset. For instance, 47 out of the 90 species that were used in the species-specific models stem from Trinidad. Accordingly, it is not surprising that the species with the highest observed morphological divergence are from Trinidad (*Chlorophanes spiza, Dysithamnus mentalis, Myiodynastes maculatus, Myrmotherula axillaris,* and *Tolmomyias sulphurescens*). Nevertheless, some general properties of Trinidad might explain why the species with the highest divergence between island and mainland are all from this island. We hypothesize that Trinidad offers a biogeographical setting that allows island-mainland intraspecific differentiation. Ecological release resulting from species impoverishment and a reduction in predators on the island compared to mainland South America could be a likely driver of morphological divergence, but not necessarily conducive to endemism. Close proximity of Trinidad to the continent (less than 15 km, the closest distance to the continent across the four islands considered here) suggests a higher chance of gene flow between island and mainland populations, thus counteracting full speciation. Still, this high level of morphological divergence calls for an integrative taxonomic approach, including molecular and song data, to clarify whether these island populations may already constitute distinct endemic species to Trinidad. Currently, only one of them, *Tolmomyias sulphurescens*, is considered an endemic subspecies of Trinidad according to Birds of the World (subspecies *berlepschi*).

## Conclusion

Our findings suggest that land-bridge islands exert similar evolutionary pressures on morphology as oceanic islands. This is evidenced by the detected combination of predictable morphological changes following the island syndrome, but also by species-specific responses. According to Alfred Russel Wallace “In islands we have the facts of distribution often presented to us in their simplest forms, along with others which become gradually more and more complex; and we are therefore able to proceed step by step in the solution of the problems they present.” (Wallace, 1880). Being more complex than oceanic islands, but still not as complex as the mainland, land-bridge islands are a promising study system to advance our knowledge on evolution and species community assembly. As their name indicates, as study systems, they are a bridge between oceanic islands and the mainland.

## Data availability

All data and code used in this study are available in the supplementary data.

## Acknowledgments

We thank the government of Equatorial Guinea for issuing research permits and INDEFOR-AP (Instituto Nacional de Desarrollo Forestal y Manejo del Sistema de Áreas Protegidas) for access to field sites and permission to capture and measure bird specimens in the Pico Basilé National Park, and Caldera de Luba Scientific Reserve. Pepijn Kamminga for support regarding the use of the ornithological collection of Naturalis Biodiversity Center. Catherine Sheard for advice on performing measurements of museum specimens.

