## Supplementary Table 3, 5, 6. 7 and Supplementary figure 1, 2, 3, 4 and 5 for "Morphological shifts consistent with the island syndrome in land-bridge island birds"

### Supplementary tables and figures

**Table S3.** Summary statistics for the ten morphological traits measured in this study. Coverage refers to the percentage of specimens for which there is data for this trait.

| Trait | Sample size | Coverage (%) | Min (mm) | Max (mm) | Mean (mm) |
|---|---|---|---|---|---|
| **Beak length culmen** | 4,683 | 67.7 | 5.2 | 59.9 | 16.6 |
| **Beak length nares** | 3,278 | 47.4 | 3.3 | 38.9 | 11.4 |
| **Beak depth** | 4,317 | 62.4 | 1.9 | 21.4 | 5.4 |
| **Beak width** | 3,580 | 51.8 | 1.9 | 15.3 | 5.2 |
| **Tarsus length** | 5,183 | 74.9 | 5.6 | 52.8 | 20.1 |
| **Primary wing length** | 6,838 | 98.9 | 36.0 | 286.0 | 74.4 |
| **Secondary wing length** | 3,549 | 51.3 | 34.1 | 189.9 | 61.5 |
| **Kipp's distance** | 3,549 | 51.3 | 1.0 | 111.1 | 14.1 |
| **Hand-wing index**[1] | 3,549 | 51.3 | 2.2 | 54.7 | 17.6 |
| **Tail length** | 6,437 | 93.1 | 18.0 | 205.0 | 59.6 |

**[1] Hand-wing index values for min, max and mean are not in mm, they are calculated as the proportion of Kipp's distance to the primary wing length times 100.**

**Table S5.** Summary of the variables used in the model.

| Element | Type | Role in model | Description |
|---|---|---|---|
| **Trait** | Continuous | Response variable | Beak length culmen<br>Beak depth<br>Beak width<br>Tarsus length<br>Primary wing length<br>Hand-wing index<br>Tail length<br>Principal Component 1 (PC1)<br>Principal Component 2 (PC2) |
| **Category Region** | Factor | Main effect | Mainland<br>Land-Bridge Island (LBI) |
| **Species** | Factor | Group-level effect | 185 bird species |
| **Data type** | Factor | Group-level effect | Field Bioko<br>Field AVONET<br>Museum AVONET<br>Museum Hoepel |

#### Passerine-wide model settings

As described in the main text, for each trait we ran a Bayesian model. As sample size and trait values changed among traits, each model had different priors and model settings. These are listed in the tables below.

**Table S6.** Summary of the prior distributions applied for each trait model. Priors were informed by data summaries, model expectations and prior knowledge as explained in our hypothesis. The intercept values reflect the expected mean for that trait on a log scale for mainland specimens. Fixed effect (category region) reflects the expected shift for trait for land-bridge island specimens. Random effect (species & data type) reflects the expected variability across species and data types. Residual priors shows the expected remaining variation for individuals

| Trait/PC | Intercept prior | Fixed effect prior (Category region) | Random effect prior (Species) | Random effect prior (Data type) | Residuals prior |
|---|---|---|---|---|---|
| **Beak length culmen** | normal(2.7, 0.1) | normal(0, 0.025) | student-t(3, 0, 1) | exponential(2) | normal(0, 0.4) |
| **Beak depth** | normal(1.61, 0.15) | normal(0, 0.2) | student-t(3, 0, 1) | exponential(2) | normal(0, 0.4) |
| **Beak width** | normal(1.6, 0.1) | normal(0.01, 0.02) | student-t(3, 0, 1) | exponential(3) | normal(0, 0.2) |
| **Tarsus length** | normal(2.97, 0.1) | normal(0, 0.025) | student-t(3, 0, 1) | exponential(2) | normal(0, 0.3) |
| **Primary wing length** | normal(4.26, 0.1) | normal(0, 0.02) | student-t(3, 0, 1) | exponential(3) | normal(0, 0.35) |
| **Hand-wing index** | normal(2.8, 0.1) | normal(-0.02, 0.02) | student-t(3, 0, 1) | exponential(2) | normal(0, 0.4) |
| **Tail length** | normal(4, 0.1) | normal(0, 0.02) | student-t(3, 0, 1) | exponential(1.5) | normal(0, 0.4) |
| **PC1** | normal(0, 0.5) | normal(0, 0.1) | student-t(3, 0, 1) | exponential(2) | normal(0, 1) |
| **PC2** | normal(0, 0.4) | normal(0, 0.1) | student-t(3, 0, 1) | exponential(2) | normal(0, 1) |

**Table S7.** Model fitting settings for Bayesian models for each trait or principal component. Log-transformed refers to whether the trait values were log transformed prior to analysis. Total number of MCMC-iterations and warm-ups steps are reported. adapt-delta value and maximum tree depth are specific to the brms packages. These model settings were optimized per trait to ensure complete chain convergence, as well as efficiency of sampling.

| Trait/PC | Log-transformed | Iterations | Warmup | adapt_delta | max_treedepth |
|---|---|---|---|---|---|
| **Beak length culmen** | Yes | 20000 | 10000 | 0.9995 | 15 |
| **Beak depth** | Yes | 40000 | 20000 | 0.9999 | 15 |
| **Beak width** | Yes | 40000 | 20000 | 0.9999 | 15 |
| **Tarsus length** | Yes | 20000 | 10000 | 0.9995 | 15 |
| **Primary wing length** | Yes | 60000 | 30000 | 0.9999 | 15 |
| **Hand-wing index** | Yes | 30000 | 15000 | 0.9995 | 15 |
| **Tail length** | Yes | 50000 | 25000 | 0.9995 | 15 |
| **PC1** | No | 60000 | 30000 | 0.9995 | 15 |
| **PC2** | No | 50000 | 25000 | 0.9995 | 15 |

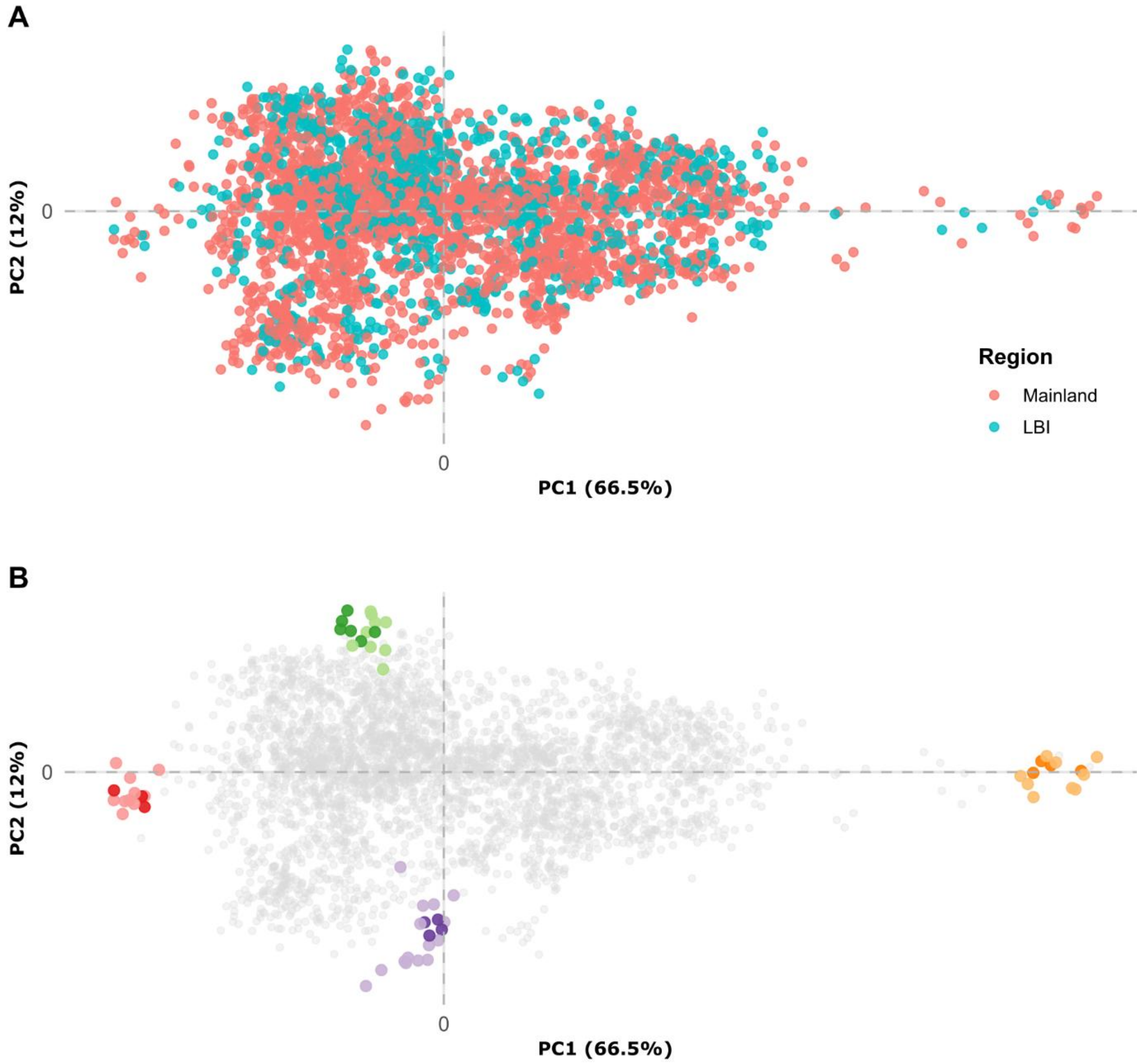


**Figure S1.** Principal component analyses of morphological data on all specimens with complete data for five traits (beak length, beak depth, tarsus length, primary wing length and tarsus length), comprising 3,301 individual specimens from 185 species. (A) PCA with each specimen coloured by region, either mainland or land-bridge island (LBI). (B) same as A, but with four species highlighted to show four distinct morphologies along morphological trait space, with darker shade indicating specimens from land-bridge islands and lighter shade indicating specimens from the mainland. In green is *Ramphocaenus melanurus* (averaged-sized bird with a long, thin beak); in yellow is *Corvus splendens* (large-sized bird with an average shaped beak); in purple is *Sporophila angolensis* (averaged-sized bird with a short, thick beak) and in red is *Dicaeum ignipectus* (small-sized bird with an average shaped beak).

#### Species-specific models output

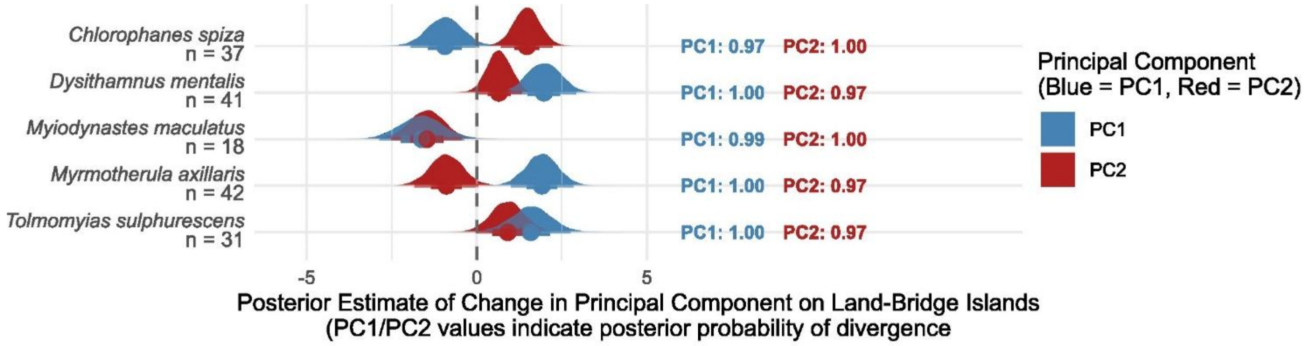


**Figure S2:** Species-specific models. Posterior estimates of morphological divergence for the species classified in the divergence category "differentiation in both" (n = 5). For each species, posterior estimates of divergence for the first two principal components are shown (PC1 = blue, PC2 = red). Dashed line indicates no difference with the mainland, and horizontal bars show 66% and 95% credible intervals. PC1 and PC2 values represent the posterior probability that the estimated effect for that principal component differs from zero, which is the mainland value.

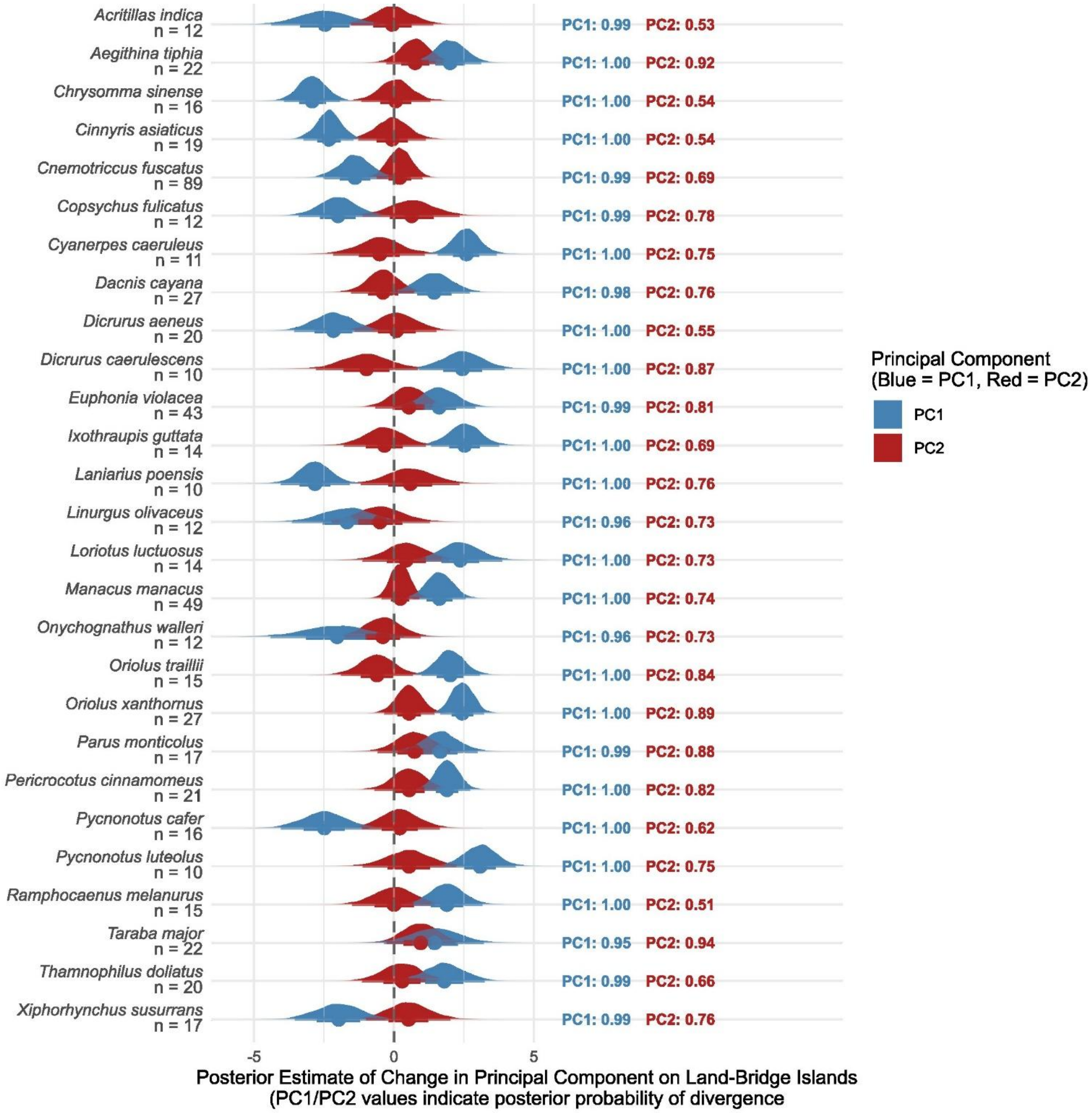


**Figure S3:** Species-specific models. Posterior estimates of morphological divergence for the species classified in the divergence category "differentiation in PC1 only" (n = 27). For each species, posterior estimates of divergence for the first two principal components are shown (PC1 = blue, PC2 = red). Dashed line indicates no difference with the mainland, and horizontal bars show 66% and 95% credible intervals. PC1 and PC2 values represent the posterior probability that the estimated effect for that principal component differs from zero, which is the mainland value.

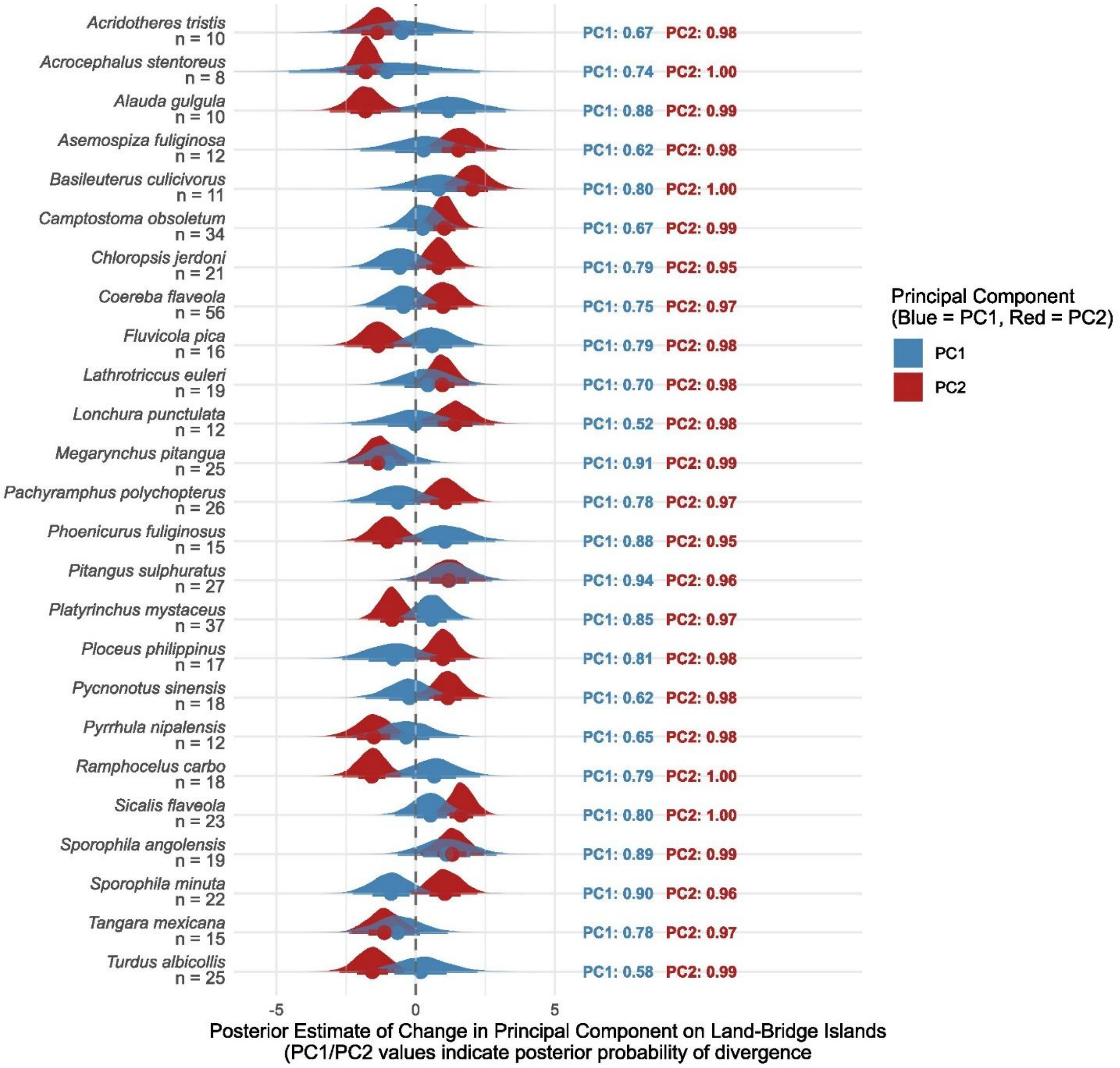


**Figure S4:** Species-specific models. Posterior estimates of morphological divergence for the species classified in the divergence category "differentiation in PC2 only" (n = 25). For each species, posterior estimates of divergence for the first two principal components are shown (PC1 = blue, PC2 = red). Dashed line indicates no difference with the mainland, and horizontal bars show 66% and 95% credible intervals. PC1 and PC2 values represent the posterior probability that the estimated effect for that principal component differs from zero, which is the mainland value.

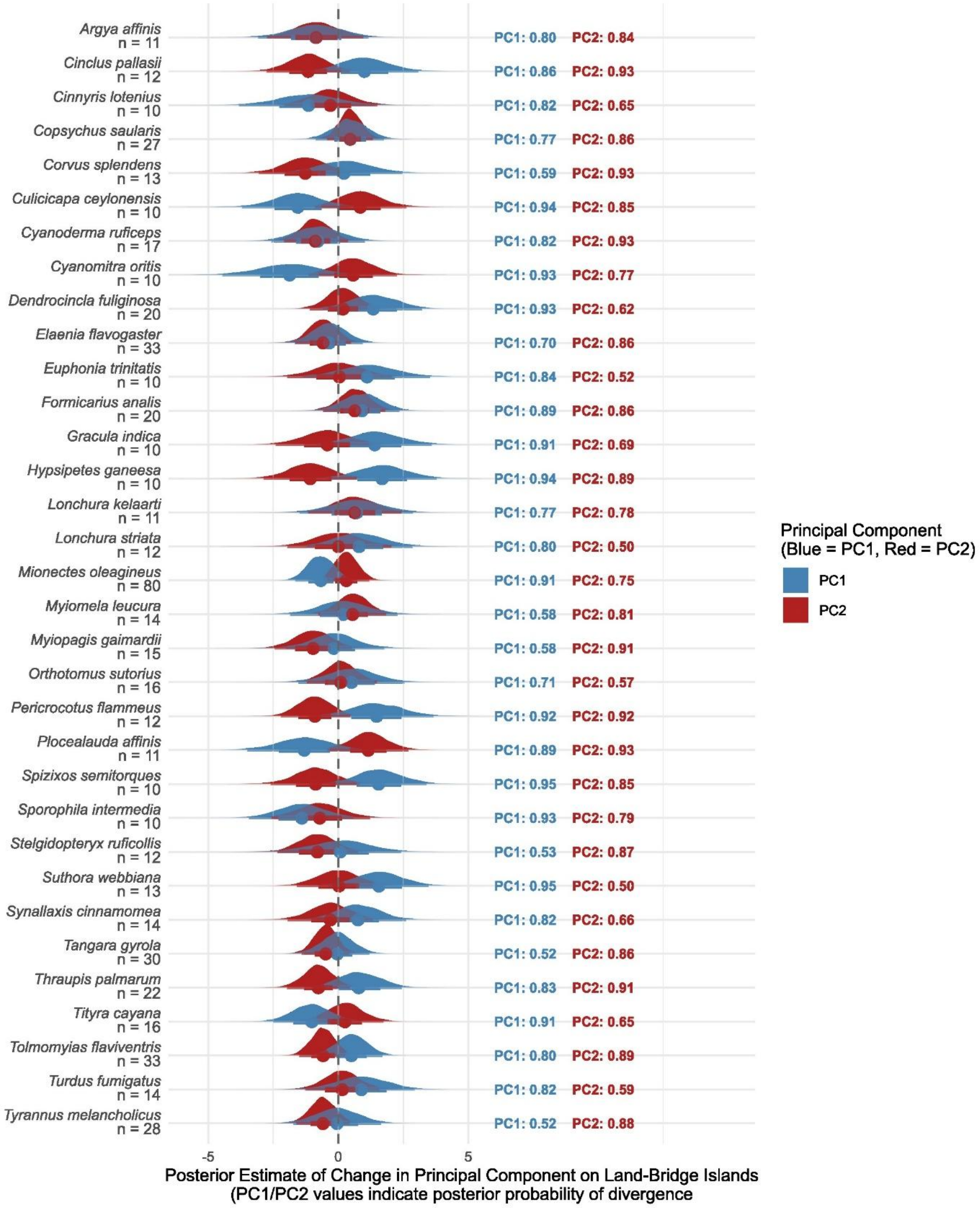


**Figure S5:** Species-specific models. Posterior estimates of morphological divergence for the species classified in the divergence category "differentiation in none" (n = 33). For each species, posterior estimates of divergence for the first two principal components are shown (PC1 = blue, PC2 = red). Dashed line indicates no difference with the mainland, and horizontal bars show 66% and 95% credible intervals. PC1 and PC2 values represent the posterior probability that the estimated effect for that principal component differs from zero, which is the mainland value.
